# Microbially conjugated bile acids differentially affect the lifecycle and pathogenesis of *Clostridioides difficile*

**DOI:** 10.64898/2026.11.05.764496

**Authors:** Sam W. Hamacher, Nitesh K. Nandwana, Noah S. Lyons, Abigail S. Gancz, Cypress E. Perkins, Brian Cheng, Emily C. Gentry, Casey M. Theriot

## Abstract

*Clostridioides difficile* is a Gram-positive bacterium that causes *C. difficile* infection (CDI), which accounts for approximately 500,000 cases and 29,000 deaths annually in the United States. In healthy adults, it has been proposed that the gut microbiota provides colonization resistance against *C. difficile* by competition through nutrient limitation and antagonism through secondary bile acid metabolism, the latter of which is mediated by the deconjugation of host-derived bile acids with bacterial bile salt hydrolases (BSH). Recently, BSHs have also shown an ability to reconjugate non-canonical (i.e., not glycine or taurine) amino acids to sterol cores, resulting in novel microbially conjugated bile acids (MCBAs). MCBAs have been associated with the resolution of recurrent CDI, though the mechanism underlying these effects are not well understood. To address this gap, we synthesized 36 MCBAs with nine amino acids (cysteine, isoleucine, leucine, phenylalanine, proline, serine, tryptophan, tyrosine, and valine) conjugated to the primary and secondary bile acids found in the human gut (CA, DCA, CDCA, and LCA) and tested their impact on various stages of *C. difficile*’s lifecycle including bacterial outgrowth, membrane integrity, sporulation, and toxin activity. We show that BSH-mediated amidation of bile acid sterol cores can either increase or decrease inhibitory effects on *C. difficile* in an amino acid-dependent manner. For example, serine MCBAs enhanced *C. difficile*’s ability to grow in otherwise inhibitory bile acid concentrations, increased spore production, and elevated toxin activity. On the other hand, tryptophan MCBAs increased inhibition. By demonstrating these variable effects, our work clarifies the molecular determinants of MCBA activity and highlights specific MCBAs as promising targets for the development of *C. difficile* microbial or molecular-based therapeutics.

**Importance:** *Clostridioides difficile* infection (CDI) remains a serious public health threat. Increasing evidence links gut microbial bile salt hydrolase (BSH) activity and the resulting microbially conjugated bile acids (MCBAs) to colonization resistance and amelioration of this disease. Although novel MCBAs continue to be identified in the human gut, their functional effects on *C. difficile* throughout its lifecycle remain largely uncharacterized. Here, we systematically evaluate how key MCBAs, including those conjugated to non-canonical amino acids for which *C. difficile* has biosynthetic auxotrophies, influence bacterial outgrowth, membrane integrity, sporulation, and toxin activity. These findings highlight the functional consequences of bile acid amidation for *C. difficile* biology and inform the rational design of novel therapeutics against CDI.

## Introduction

*Clostridioides difficile* is a Gram-positive, spore-forming bacterium that causes *C. difficile* infection (CDI), a disease characterized by symptoms ranging from mild to moderate diarrhea to severe complications, including pseudomembranous colitis and death. The Centers for Disease Control and Prevention (CDC) classifies CDI as a serious public health threat, with approximately 500,000 cases and 29,000 deaths annually in the United States (1). Although antibiotics such as vancomycin and fidaxomicin are the standard of care for CDI, up to 35% of primary CDI patients will experience recurrence (2). These recurrences are thought to occur due to the loss of colonization resistance (CR), a phenotype mediated by gut microbiota in the healthy gut (3, 4). Antibiotic exposure disrupts the composition, structure, and function of the gut microbiota, creating an environment more permissive to *C. difficile* colonization (5, 6). Although CR encompasses multiple mechanisms, two well-characterized contributors are gut microbiota-mediated modifications of the host bile acid (BA) pool and competition with *C. difficile* for ecological niches and nutrients (7, 8). Historically, these mechanisms have often been considered distinct, with BA metabolism understood primarily in terms of the production of metabolites that inhibit *C. difficile*, while nutrient competition has been studied as a separate mechanism of CR (7, 9). However, the recent discovery of novel, microbially modified bile acids raise questions about the interplay between these processes and their relative contributions to CR and disease.

Host-derived BAs are synthesized from cholesterol in the liver, conjugated to taurine or glycine, and secreted into the small intestine as conjugated primary BAs (10). Within the small intestine, widely distributed microbial enzymes known as bile salt hydrolases (BSHs) deconjugate primary BAs, releasing the unconjugated BA and free amino acid (11). BSH-mediated deconjugation acts as a gatekeeper of downstream BA metabolism because many subsequent microbial transformations require the unconjugated sterol cores of cholic acid (CA) or chenodeoxycholic acid (CDCA) (12, 13) (Figure 1A). For example, the secondary BAs deoxycholic acid (DCA) and lithocholic acid (LCA) are produced through 7α-dehydroxylation of CA and CDCA, respectively. This transformation is mediated by the bile acid-inducible (*bai*) operon, which is primarily encoded by the Class Clostridia (11, 14–16). Secondary BAs and certain derivatives, including their epimers (e.g., isoDCA, isoLCA), have demonstrated inhibition of *C. difficile* growth and virulence (17–20).

**Figure 1.**
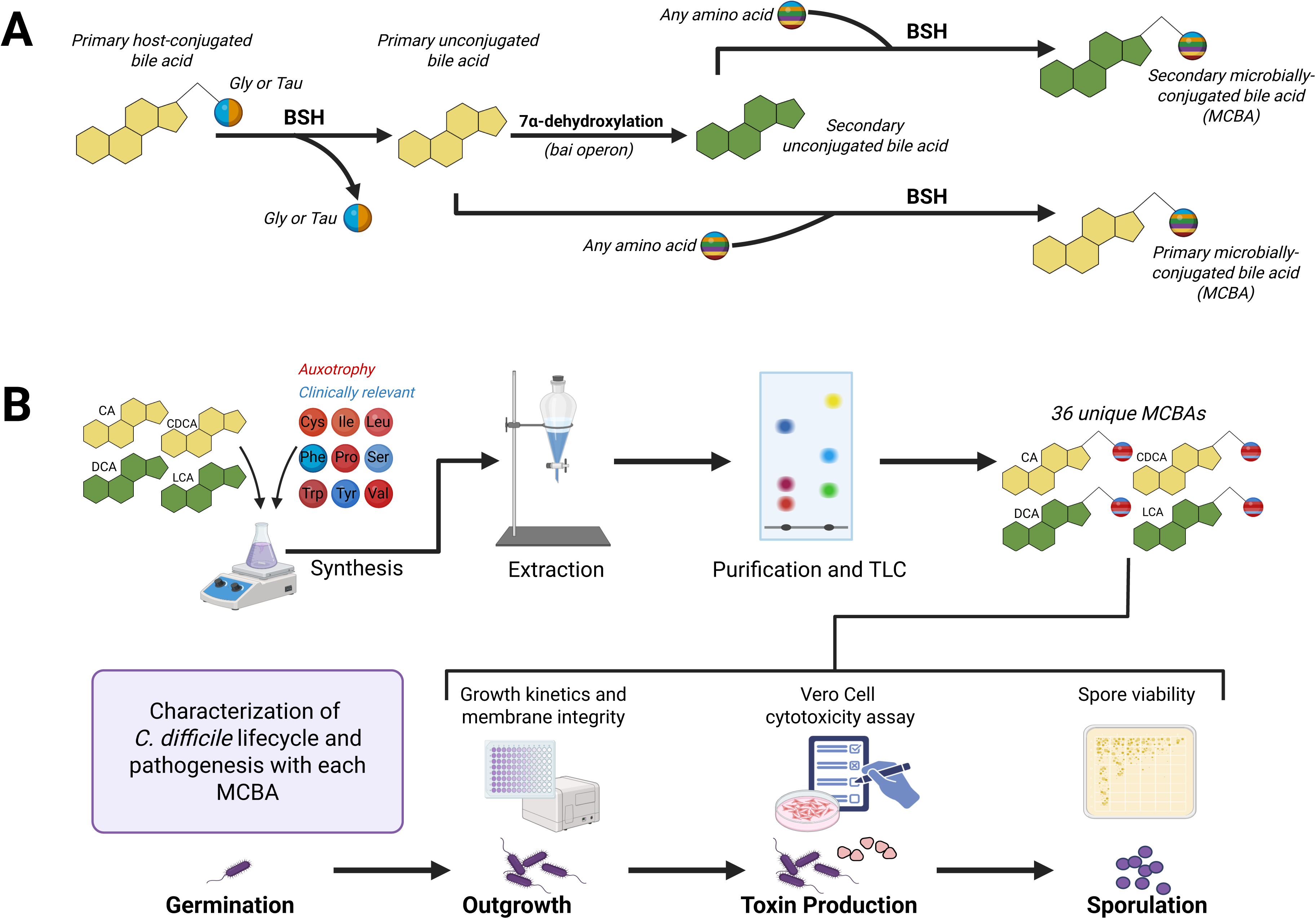
Bacterially encoded BSHs modify host produced BAs to create microbially conjugated bile acids (MCBAs). A) Overview of bile acid metabolism. Host produced primary conjugated BAs are deconjugated by BSHs that leave free primary BA cores, which can then be transformed through the *bai* operon to secondary BAs. BSHs also catalyze the MCBA formation of both primary and secondary bile acid cores, resulting in primary and secondary MCBAs. B) Schematic showing the MCBAs synthesized for this study and how they were tested in each assay with *C. difficile*. Each bile acid core has each amino acid, resulting in nine bile acids per sterol core (AA-CA, AA-DCA, AA-CDCA, AA-LCA).

Expanding the traditional repertoire of primary and secondary BAs, recent publications have identified a plethora of microbially conjugated bile acids (MCBAs) throughout the human intestinal tract at biologically relevant concentrations (21–23). Recently, it was found that BSHs are responsible for making MCBAs via a secondary acyltransferase activity, reconjugating amino acids beyond glycine or taurine onto free sterol cores. *Clostridium perfringens* BSH has broad MCBA reconjugation activity with little preference for the amino acid substrate, while *Bifidobacterium longum* BSH prefers more aromatic amino acid substrates (22, 23). In addition to amino acid substrates, BSHs with reconjugation activity show preference for sterol cores, resulting in both primary and secondary MCBAs (24, 25). Beyond reconjugation, BSHs show deconjugation activity specifically on MCBAs, as BSHs from *Lactobacillus* can deconjugate a wide pool of CA amidates (26, 27). The newly described bifunctional activity of BSHs and the production of MCBAs with diverse BA sterol cores conjugated to multiple amino acids substantially increases the known complexity of the BA pool. However, the effects of these novel metabolites on the host, the gut microbiota, and *C. difficile* remain largely undefined.

Understanding these host-pathogen effects is critical as canonical BAs are heavily involved in host signaling, including through the farnesoid X receptor (FXR) which regulates BA homeostasis and hepatic BA synthesis (28–30). The presence of MCBAs across multiple anatomical sites, with variation in their abundance according to location and time, raises the possibility that they also participate in BA signaling and influence host physiology beyond the intestines (31–33). MCBAs may also influence *C. difficile* pathogenesis. In antibiotic-treated mice, administration of a cocktail of BSHs from different *Lactobacillus* species increased the abundance of both deconjugated BAs and MCBAs and was associated with a reduced *C. difficile* burden during early infection. Among MCBAs detected *in vivo*, phenylalanine-conjugated CA (Phe-CA) and tyrosine-conjugated CA (Tyr-CA) reduced *C. difficile* spore germination in the presence of the germinant TCA and altered toxin expression *in vitro* (26). Furthermore, MCBA profiles differ between pre- and post-fecal microbiota transplantation (FMT) samples from patients with rCDI, with successful FMT associated with a shift from higher levels of primary BA-conjugated MCBAs toward secondary BA-conjugated MCBAs, such as DCA and LCA conjugates (34).

Despite these emerging findings, the biological activities of individual MCBAs remain poorly characterized, in part because these compounds are not commercially available and must be chemically synthesized. Consequently, their effects on *C. difficile* growth, membrane integrity, toxin activity, and sporulation remain poorly described. Given that both BA-mediated inhibition and amino acid competition are established contributors to CR and virulence, we sought to determine whether these mechanisms intersect through MCBA activity. Specifically, we hypothesized that conjugation to amino acids that *C. difficile* cannot synthesize (i.e., amino acids for which it has auxotrophies) would enhance the inhibitory effects of BAs on *C. difficile* relative to their unconjugated sterol cores. To investigate this, we characterized 36 MCBAs comprising nine non-canonical amino acid substrates (cysteine, isoleucine, leucine, phenylalanine, proline, serine, tryptophan, tyrosine, and valine) across four canonical BA sterol cores (Figure 1B). We evaluated their effects on *C. difficile* growth, membrane integrity, spore formation and toxin activity. By systematically evaluating the effects of these MCBAs on *C. difficile* pathogenesis, this work investigates a potential link between BA-mediated effects and nutrient competition in CR and may inform the development of novel BA-based therapeutics for CDI and other intestinal diseases.

## Results

### MCBAs with CA in the sterol core (AA-CA) are less inhibitory to *C. difficile* than cholate

Growth curves measuring the area under the curve (AUC) showed that all CA concentrations tested (1.25 mM, 2.5 mM, 5 mM, and 10 mM) affected *C. difficile* growth over 24 hr when compared to the vehicle control (DMSO, no bile acid), with 10 mM CA completely inhibiting growth (Figure 2A, Figure S1A-D). Cys-, Pro-, Ser-, and Val-CA did not inhibit *C. difficile* growth when supplemented with up to 10 mM of MCBAs. Tyr-CA had higher growth compared to the vehicle when supplemented with 10 mM (Figure S1E). Ser-CA at 2.5 mM, 5 mM, and 10 mM had a subtle decrease in growth. Leu-CA had minimal inhibition up to 5 mM and then became more inhibitory at 10 mM, which differs from CA’s stepwise decrease in AUC. Ile-CA, Phe-CA and Trp-CA inhibited *C. difficile* growth at 5 mM and above.

**Figure 2.**
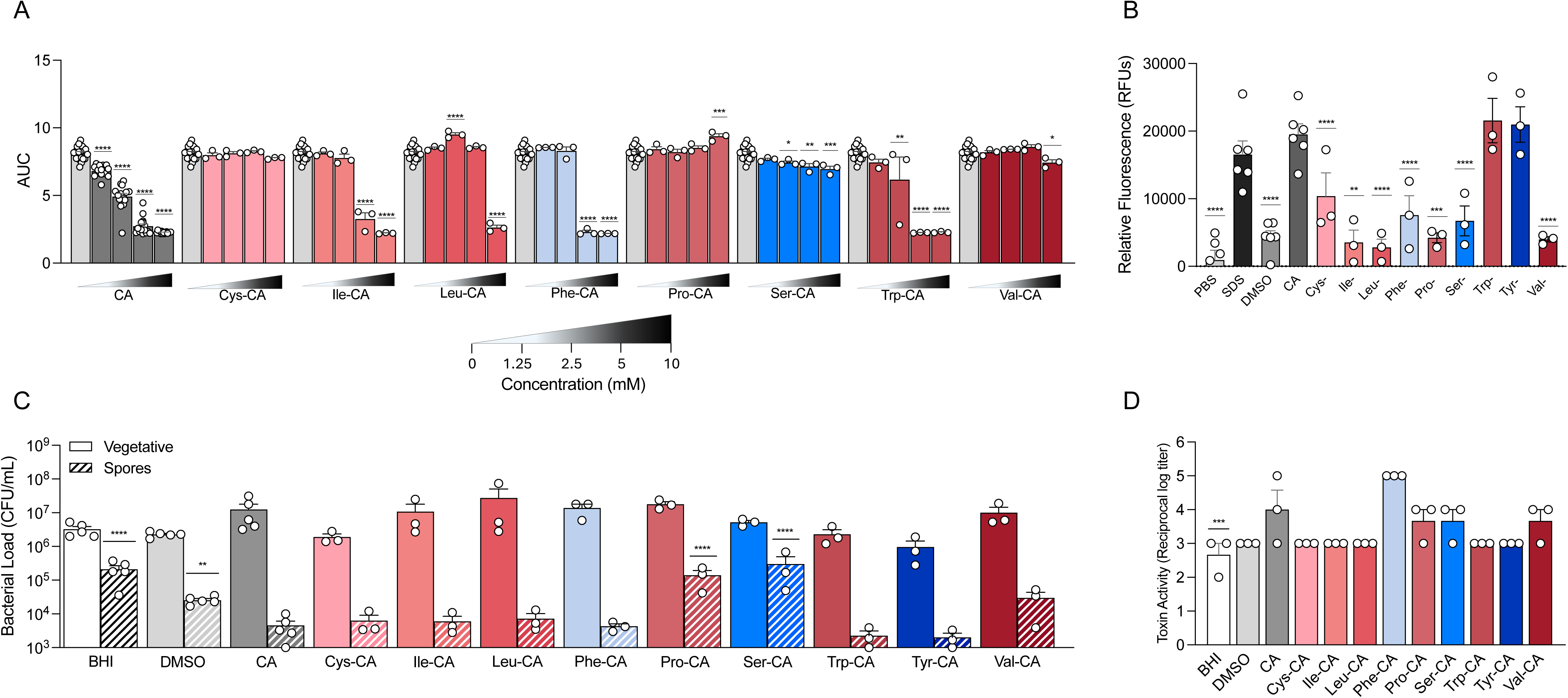
CA amidates are less inhibitory to *C. difficile* than cholate. A) *C. difficile* growth curves were done by measuring OD_600_ over 24 hr in varying concentrations of MCBAs. Area Under the Curve (AUC) is displayed. Concentrations ranged from 0 mM (2.5% DMSO, gray bar) 1.25 mM, 2.5 mM, 5 mM, and 10 mM. Each bile acid was run in triplicate, with 3 biological replicates. Significance was determined by ordinary one-way ANOVA with Dunnett’s posttest comparing each concentration to the 0 mM condition. B) Membrane integrity assay using a propidium iodide stain of *C. difficile* cells after incubation in 2.5 mM of AA-CAs, normalized to the background fluorescence. Significance was determined by ordinary one-way ANOVA with Dunnett’s posttest comparing each MCBA to CA. C) Enumeration of vegetative cells (solid bars) and spores (striped bars) after incubation in 2.5 mM MCBAs for 72 hr. Significance was determined by ordinary one-way ANOVA with Dunnett’s posttest comparing each MCBA to CA. D) *C. difficile* toxin activity via a Vero Cell cytotoxicity assay after incubation in AA-CAs for 72 hr, significance was determined by ordinary one-way ANOVA with Dunnett’s posttest comparing each MCBA to CA. * = p< 0.05, ** = p<0.01, ***= p<0.001, **** = p< 0.0001.

In membrane integrity assays at 2.5 mM, CA had more membrane disruption than both the PBS and vehicle controls (DMSO) and had similar levels of disruption compared to the positive control SDS (Figure 2B). The addition of 2.5 mM Cys-, Ile-, Leu-, Phe-, Pro-, Ser- and Val-CA reduced its ability to disrupt the membrane when compared to CA. Trp-CA and Tyr-CA mirrored CA in that it was able to disrupt the membrane, maintaining a high level of fluorescence. Overall, amidates of CA were able to decrease membrane disruption compared to the sterol core CA.

We next sought to determine how different amidates of CA affect vegetative cell growth and spore formation over a 72 hr period. Compared to both BHI and DMSO controls, supplementation of 2.5 mM CA reduced spores recovered on TBHI medium at 72 hr without affecting overall vegetative cell growth (Figure 2C). Trp-CA and Tyr-CA increased vegetative cell growth at 24 hr only, and all amidates exhibited similar growth and spore formation compared to CA at 48 hr. (Figure S1F-G). Cys-, Ile-, Leu-, Phe-, Trp-, Tyr-, and Val-CA also reduced spore formation, similar to CA alone with no changes in vegetative cell growth (Figure 2C). Pro- and Ser-CA did not decrease spore formation, restoring spores to similar levels as the controls. Finally, supplementation of 2.5 mM CA increased toxin activity at 72 hr compared to BHI alone but not in the DMSO control. CA amidates had similar levels of toxin activity compared to CA alone at both 24 hr and 72 hr (Figure 2D, Figure S1H).

### MCBAs with DCA in the sterol core (AA-DCA) rescue DCA inhibition of *C. difficile* growth, spore formation, and toxin activity

DCA amidates exhibited different effects compared to CA amidates. DCA alone is more inhibitory than CA and has a lower MIC at 1.2 mM, and thus we predicted that AA-DCAs would have increased inhibitory effects compared to CA amidates. As the concentration of DCA increased from 0.075 mM, 0.3 mM, 0.6 mM, and 1.2 mM it decreased *C. difficile* growth compared to the vehicle control (DMSO, no bile acid), with total inhibition at 1.2 mM (Figure 3A, Figure S2A-D). Interestingly, Cys-, Pro-, Ser-, and Tyr-DCA reversed this effect. Cys-, Pro-, and Ser-DCA had a small decrease in AUC but still had robust growth at 1.2 mM, and Tyr-DCA had an increase in growth at 1.2 mM compared to the vehicle. Ile-, Leu-, and Phe-DCA had a more pronounced decrease in AUC at lower concentrations, becoming inhibitory at 0.6 mM. Trp-DCA was inhibitory at 0.3 mM, well below the 1.2 mM MIC of DCA. Lastly, Val-DCA followed a similar trend to DCA alone, except at 0.3 mM where there was no difference in growth compared to the vehicle and then a sharp decrease in 0.6 mM.

**Figure 3.**
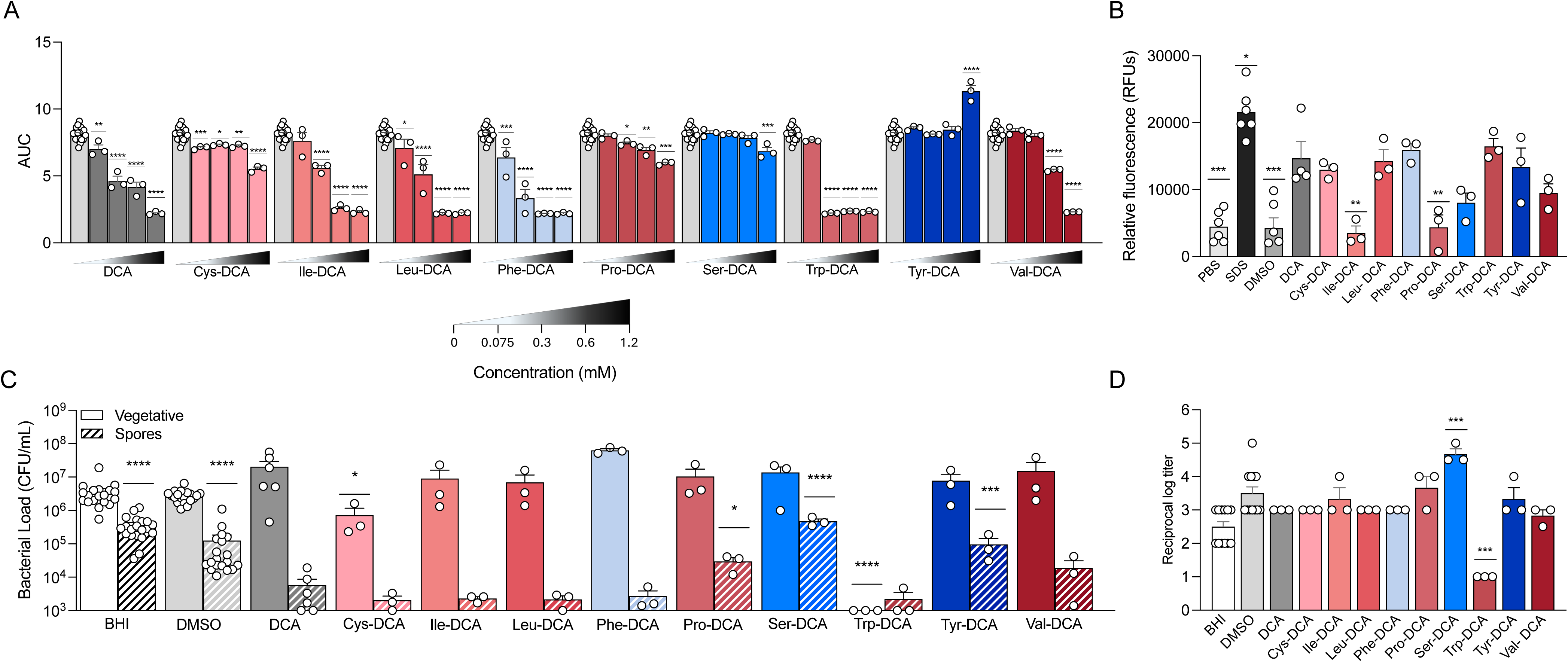
DCA amidates reverse DCA inhibition of *C. difficile* growth, spore formation, and toxin activity. A) *C. difficile* growth curves were done by measuring OD_600_ over 24 hr in varying concentrations of MCBAs. Area Under the Curve (AUC) is displayed. Concentrations ranged from 0 mM (2.5% DMSO, gray bar) 0.075 mM, 0.3 mM, 0.6 mM, and 1.2 mM. Each bile acid was run in triplicate, with 3 biological replicates. Significance was determined by ordinary one-way ANOVA with Dunnett’s posttest comparing each concentration to the 0 mM condition. B) Membrane integrity assay using a propidium iodide stain of *C. difficile* cells after incubation in 0.3 mM of AA-DCAs, normalized to the background fluorescence. Significance was determined by ordinary one-way ANOVA with Dunnett’s posttest comparing each MCBA to DCA. C) Enumeration of vegetative cells (solid bars) and spores (striped bars) after incubation in MCBAs for 72 hr. Significance was determined by ordinary one-way ANOVA with Dunnett’s posttest comparing each MCBA to DCA D) *C. difficile* toxin activity via a Vero Cell cytotoxicity assay after incubation in AA-DCAs for 72 hr, significance was determined by ordinary one-way ANOVA with Dunnett’s posttest comparing each MCBA to DCA. * = p< 0.05, ** = p<0.01, ***= p<0.001, **** = p< 0.0001.

In membrane integrity assays at 0.3 mM, DCA disrupted the membrane more than the PBS and DMSO negative controls but less than the SDS positive control (Figure 3B). When comparing the DCA amidates to DCA alone, only Ile-DCA and Pro-DCA decreased membrane disruption compared to DCA, despite Ile-DCA being more inhibitory to growth than DCA and Pro-DCA increasing growth. The other AA-DCAs did not alter DCA mediated effects on membrane integrity although Ser-DCA had subtle, but not statistically significant, disruption to membrane disruption.

Next, we examined the effects of DCA amidates on vegetative cell growth and spore formation when incubated in 0.3 mM AA-DCAs for a 72 hr period. After both 48 and 72 hr, DCA reduced spore recovery compared to BHI and the DMSO controls (Figure 3C, Figure S2F). Most of the DCA amidates did not affect spore formation and looked similar to DCA, however Pro-, Ser-, and Tyr-DCA increased spores compared to DCA at 72 hr (Figure 3C, Figure S2E-F). Interestingly, Pro-DCA did not have a significant increase in sporulation until 72 hr, whereas Ser- and Tyr-DCA had higher spore counts at 48 hr (Figure S2F). Cys- and Trp-DCA decreased vegetative cells at 72 hr. Lastly, when examining toxin activity 0.3 mM DCA supplementation did not change the overall toxin activity compared to the BHI control but had lower toxin activity than the DMSO control at 24 and 72 hr. (Figure 3D, Figure S2G). In line with the spore formation assay, most DCA amidates had a similar effect on toxin activity compared to DCA. Trp-DCA had less toxin activity, however at 72 hr of growth there were no detectable vegetative cells, which explains low toxin activity. Of the DCA amidates that increased spore formation, only Ser-DCA increased toxin activity at 72 hr while Pro-DCA and Tyr-DCA did not. Interestingly, at 24 hr Pro- and Tyr-DCA had increased toxin activity while Ser-DCA did not (Figure S2G).

### MCBAs with CDCA in the sterol core (AA-CDCA) affect *C. difficile* differently

Although CDCA is a primary BA, it alone is inhibitory to *C. difficile*, with an MIC at 1.2 mM similar to DCA, and it has been shown to decrease germination of *C. difficile* spores in the presence of the germinant TCA (35). We predicted that it would maintain its inhibitory effects when conjugated, and that CDCA amidates would mimic patterns seen in DCA more than those seen in CA.

CDCA was tested at the same concentrations as DCA, and showed inhibition at 1.2 mM (Figure 4A, Figure S3A-D). Ile-, Pro-, and Val-CDCA remained similar to CDCA alone (Figure 4A). Phe- and Trp-CDCA were both inhibitory at 0.3 mM, with increased inhibition compared to CDCA. Lastly, Ser- and Tyr-CDCA did not inhibit *C. difficile* with all tested MCBAs, showing robust growth in all conditions. Cys-CDCA inhibited *C. difficile* growth only at the 1.2 mM concentration.

**Figure 4.**
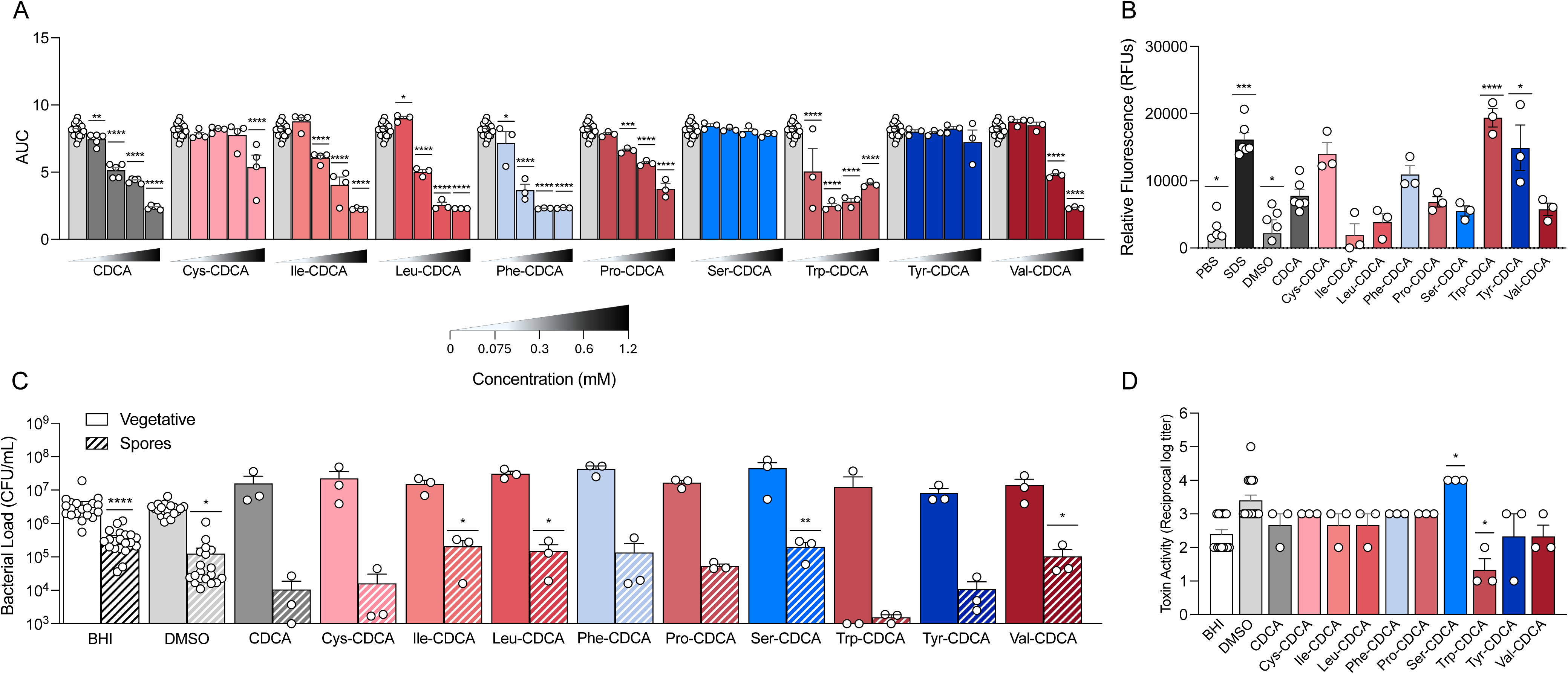
CDCA amidates affect *C. difficile* differently. A) *C. difficile* growth curves were done by measuring OD_600_ over 24 hr in varying concentrations of MCBAs. Area Under the Curve (AUC) is displayed. Concentrations ranged from 0 mM (2.5% DMSO, gray bar) 0.075 mM, 0.3 mM, 0.6 mM, and 1.2 mM. Each bile acid was run in triplicate, with 3 biological replicates. Significance was determined by ordinary one-way ANOVA with Dunnett’s posttest comparing each concentration to the 0 mM condition. B) Membrane integrity assay using a propidium iodide stain of *C. difficile* cells after incubation in 0.3 mM of AA-CDCAs, normalized to the background fluorescence. Significance was determined by ordinary one-way ANOVA with Dunnett’s posttest comparing each MCBA to CDCA. C) Enumeration of vegetative cells (solid bars) and spores (striped bars) after incubation in MCBAs for 72 hr. Significance was determined by ordinary one-way ANOVA with Dunnett’s posttest comparing each MCBA to CDCA. D) *C. difficile* toxin activity via a Vero Cell cytotoxicity assay after incubation in AA-CDCAs for 72 hr, significance was determined by ordinary one-way ANOVA with Dunnett’s posttest comparing each MCBA to CDCA. * = p< 0.05, ** = p<0.01, ***= p<0.001, **** = p< 0.0001.

In membrane integrity assays performed at 0.3 mM, CDCA alone increased the disruption of the membrane compared to the negative controls (PBS and DMSO) but less than the SDS control (Figure 4B). Interestingly, Trp- and Tyr-CDCA, an inhibitory and growth enhancing MCBA respectively, had increased membrane disruption compared to CDCA. Most CDCA amidates, however, did not change membrane disrupting effects mediated by CDCA.

AA-CDCAs at 0.3 mM at 72 hr had minor effects on *C. difficile* spore formation. After 72 hr, CDCA had less spores than the BHI control and the DMSO vehicle control (Figure 4C). Of the CDCA amidates, Ile-, Leu-, Ser-, and Val-CDCA had increased spore formation compared to CDCA, ameliorating the effects of the sterol core on spore formation at 72 hr. However, Ile-and Leu-CDCA had high spore counts at all time points whereas Ser-CDCA only had increased spores at 72 hr (Figure 4C, Figure S3E-F).

After incubation in CDCA alone for 72 hr, *C. difficile* toxin activity measured by a Vero Cell cytotoxicity assay did not change compared to the BHI and vehicle controls (Figure 4D, Figure S3G). Ser-CDCA increased toxin activity at 72 hr, while Trp-CDCA decreased toxin activity. The reduction in toxin activity by Trp-CDCA, however, may be due to poor growth over the course of 72 hr, with two of the conditions being below the limit of detectable CFUs at 72 hr (Figure 4C). Cys-, Ile-, Leu-, Phe-, Pro-, Tyr-, and Val-CDCA did not alter toxin activity compared to their sterol core CDCA at either 24 or 72 hr (Figure S3G).

### MCBAs with LCA in the sterol core (AA-LCA) are more inhibitory to *C. difficile* than LCA

Due to solubility issues, measuring optical density is not viable in the presence of LCA and its amidates. To address this, MIC assays were adapted from (23), measuring the growth of *C. difficile* in LCA at 0.15 mM, 0.3 mM, 0.6 mM, and 1.2 mM via bacterial enumeration over a 24 hr period relative to the inoculum at the start of the experiment (approximately 1×10^5^ CFU/mL). LCA showed inhibition at 1.2 mM, not growing above the inoculum dose (Figure 5A). Many of the LCA amidates were much more inhibitory compared to LCA alone, with five of the nine LCA amidates being more inhibitory than LCA (Ile, Leu, Phe, Pro, Ser, and Val), many requiring additional lower concentrations of 0.01875 mM, 0.0375 mM, and 0.075 mM tested. Of these inhibitory LCA amidates, Val-LCA had the lowest MIC at 0.0375 mM. Interestingly, Pro-LCA and Ser-LCA were inhibitory, which did not follow the trends of other sterol cores with MICs of about 0.075 mM and 0.6 mM respectively. Cys- and Tyr-LCA were the two amidates that did not inhibit growth.

**Figure 5.**
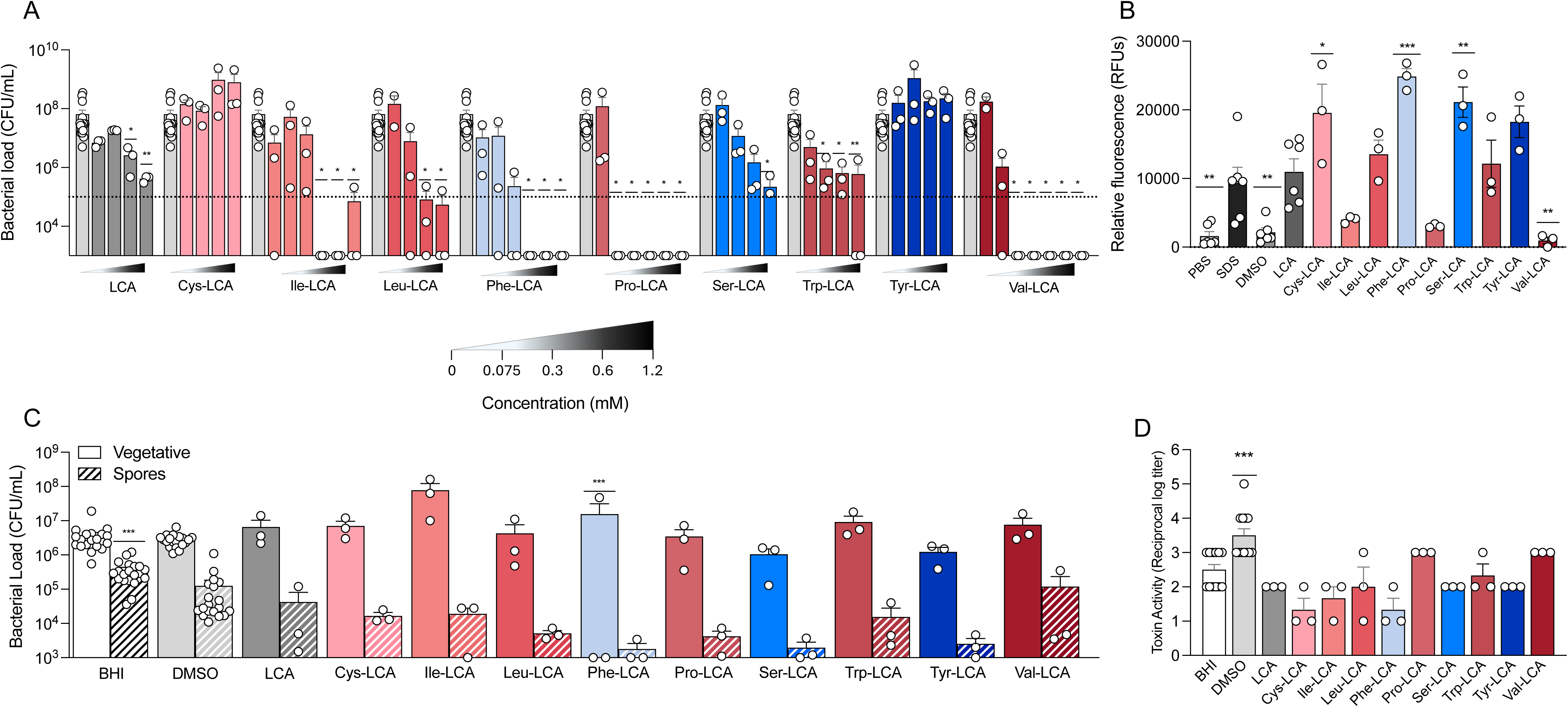
LCA amidates are more inhibitory to *C. difficile* than LCA. A) MIC Assay of bacterial enumeration of *C. difficile* after incubation in increasing concentrations of MCBAs. Concentrations ranged from 0 mM (2.5% DMSO), 0.01875 mM, 0.0375 mM, 0.075 mM, 0.15 mM, 0.3 mM, 0.6 mM, and 1.2 mM. Inhibition was determined if the enumeration after 24 hr did not exceed the inoculation (dotted line). Significance determined by Kruskal Wallis, * = p< 0.05, ** = p<0.01. B) Membrane integrity assay using a propidium iodide stain of *C. difficile* cells after incubation in 0.3 mM of AA-LCAs, normalized to the background fluorescence. Significance was determined by ordinary one-way ANOVA with Dunnett’s posttest comparing each MCBA to LCA. C) Enumeration of vegetative cells (solid bars) and spores (striped bars) after incubation in MCBAs for 72 hr. Significance was determined by ordinary one-way ANOVA with Dunnett’s posttest comparing each MCBA to LCA D) *C. difficile* toxin activity via a Vero Cell cytotoxicity assay after incubation in AA-LCAs for 72 hr, significance was determined by ordinary one-way ANOVA with Dunnett’s posttest comparing each MCBA to LCA. * = p< 0.05, ** = p<0.01, ***= p<0.001, **** = p< 0.0001.

Membrane integrity did not appear to be a major factor in the increase in growth inhibition (Figure 5B). LCA increased disruption of the membrane compared to the PBS and vehicle control and had approximately the same amount of disruption as SDS control at 0.3 mM. Cys-LCA increased membrane disruption compared to LCA, despite being less inhibitory, and Val-LCA significantly decreased membrane disruption while being more inhibitory. Phe- and Ser-LCA also increased membrane disruption compared to LCA.

We next examined effects of LCA amidates on spore formation over a 72 hr period. Due to the lower MICs of LCA amidates, we incubated *C. difficile* at a quarter of the MIC of the LCA amidate to ensure cell growth. After 72 hr LCA had a decrease in spore formation compared to the BHI control but did not differ from the vehicle control (Figure 5C). Interestingly, none of the LCA amidates altered spore formation in comparison to LCA at any time point (Figure 5C, S4A-B). Similarly, toxin activity at 72 hr revealed increased toxin activity in the DMSO control compared to LCA, but no LCA amidates altered toxin activity at 72 hr, however at 24 hr Ile- and Phe-LCA had less toxin activity (Figure 5D, Figure S4C).

## Discussion

Here we characterized 36 MCBAs comprising nine non-canonical amino acids (cysteine, isoleucine, leucine, phenylalanine, proline, serine, tryptophan, tyrosine, and valine) containing primary and secondary BA sterol cores and evaluated their effects on *C. difficile* growth, membrane integrity, sporulation, and toxin activity. Amidation with non-canonical amino acids altered the effects of unconjugated BAs, with outcomes dependent on both the amino acid and sterol core. Trp MCBAs consistently enhanced growth inhibition across all four BA cores, whereas some cysteine, proline, serine, and tyrosine MCBAs attenuated BA-mediated inhibition and promoted growth. Membrane disruption did not consistently correspond to growth inhibition, suggesting that additional mechanisms contribute to MCBA activity. MCBAs also differentially affected sporulation and toxin activity. Serine MCBAs consistently attenuated BA-mediated inhibition of sporulation and increased toxin activity, whereas several proline and tyrosine MCBAs increased spore formation without a corresponding increase in toxin activity. Despite testing a range of non-canonical amino acids, we did not observe that effects on *C. difficile*’s lifecycle and pathogenesis were linked to innate biochemical properties including hydrophobicity and aromaticity.

MCBAs were able to alter *C. difficile* growth kinetics. Tryptophan conjugates across all BA cores increased inhibition, while others like proline, serine, tyrosine, and in some cases, cysteine increased inhibition and allowed for growth. Our findings are consistent with clinical observations linking MCBA composition to CDI outcomes. Following successful FMT in patients with rCDI, tryptophan MCBAs and Val-LCA increased whereas serine MCBAs decreased (34) These changes align with the inhibitory effects of tryptophan MCBAs and the growth-promoting effects of serine MCBAs observed here (34). Similarly, a previous study in antibiotic-treated mice found that BSH treatment was associated with reduced *C. difficile* burden and increased Trp-CA and Phe-CA but also increased serine and tyrosine MCBAs (26). These findings highlight the complexity of interpreting changes in individual MCBAs *in vivo*. Although these associations do not establish a causal role for MCBAs in the prevention of *C. difficile* infection via CR, they do support further investigation into the contributions of individual MCBAs.

Effects on membrane integrity were not linked to growth inhibition in most cases, as isoleucine and proline had decreased membrane damage across the sterol cores, and tyrosine increased membrane damage despite being typically non-inhibitory. Notably, the amino acids with aromatic side chains (Phe, Trp, and Tyr) (Figure 6) consistently had higher membrane disruption, indicating that the aromatic benzene may increase the ability of BAs to enter and disrupt bacterial membranes. Across the BA cores, however, polar and non-polar amino acids did not show any trends, indicating that membrane damage may be determined more by the BA core more so than the amino acid (Figure 6). Membrane integrity is a large player in the inhibition of BAs, as their ability to embed or cross the bacterial membrane is important to their ability to disrupt cellular processes and cause membrane damage (26, 36). Sufficient concentrations of BAs that cause membrane damage would cause an inhospitable environment for *C. difficile*, although MCBAs seem to act independently of membrane damage in some cases. For example, Ile-DCA reduced membrane damage compared to DCA but was more inhibitory than DCA, and Tyr-DCA had increased damage but was not inhibitory. BAs also impact certain cellular processes, such as expression of flagellar genes like *fliC*, so it is likely that MCBAs’ mechanism of inhibition or growth promotion is less focused on membrane integrity and is more focused on their ability to alter bacterial processes (36). Looking forward, examining how MCBAs affect *C. difficile* gene expression may shed further light on their effects on growth and pathogenesis.

**Figure 6.**
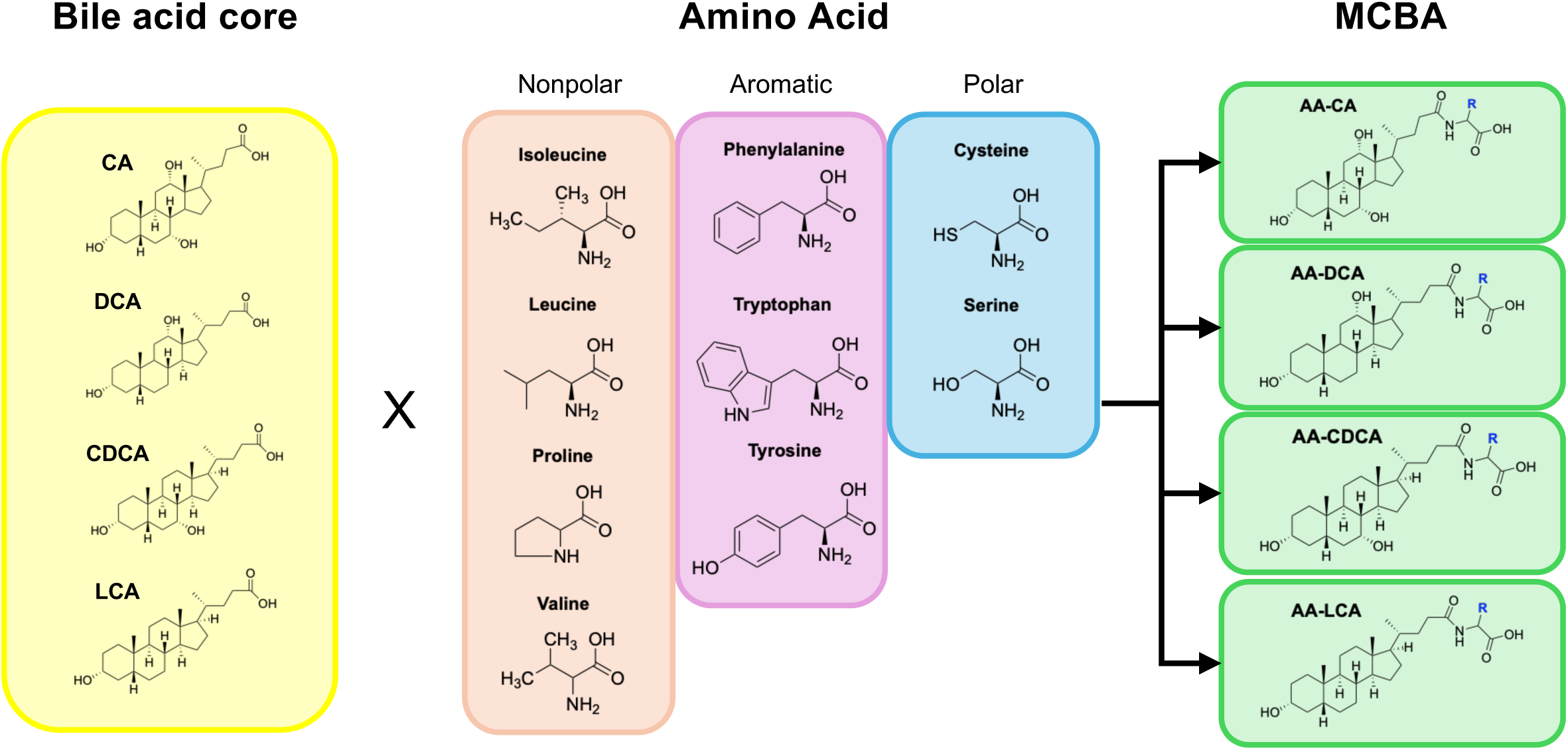
Structures of bile acids and amino acids important for MCBA creation. Structures of the 4 BA cores used in this study, and the structure of the 9 amino acids conjugated to them. Amino acid structures range from nonpolar, polar, and aromatic residues.

Amidation also changed viable spore formation; serine conjugates relieved the inhibitory effects of BAs on sporulation, as well as proline and tyrosine when conjugated to CA and DCA. Outside of serine, however, there were no other amino acids that increased sporulation across the BA cores; Pro-CA and Pro-DCA both increased sporulation, but only Tyr-DCA increased the number of spores. While there are studies that look at the effects of BAs on spore germination since TCA is a germinant, inhibition of sporulation itself is largely focused on antibiotic mediated inhibition (37–39). BAs do impact spore formation, and when considering that spores are the primary mode of transmission of *C. difficile* among patients, understanding how MCBA composition may increase or decrease transmission may be an avenue for prevention. Looking at the changes in the MCBA pool in patients post-FMT, the decreases in serine MCBAs may also correspond to less spore shedding during recovery as the microbiome restores CR (34).

Lastly, only serine conjugates increased toxin activity. While some MCBAs decreased toxin activity, the decrease was caused by lack of vegetative cell growth rather than a BA-mediated effect. Secondary BAs can reduce toxin activity *in vitro* and *in vivo,* and the amelioration of this by serine MCBAs underlies how amino acids change BA mediated effects on toxin to allow *C. difficile* pathogenesis. Also of note is Pro-CA, Pro-DCA, and Tyr-DCA’s lack of toxin increase in these assays. Toxin expression is tightly controlled by a variety of regulatory networks, including sporulation regulators *spo0A and sigH* (40, 41). These MCBAs increased spore formation *in vitro*, but did not increase toxin activity, indicating that they may affect toxin activity in a manner that represses toxin activity through other mechanisms. Considering that LCA itself can readily bind toxin, it could be that some MCBAs tested can inhibit toxin through a similar mechanism (42, 43). Examining additional aspects of toxin production, such as gene expression and toxin binding kinetics, may shed light on how these MCBAs affect toxin.

Finally, this study has several limitations. First, our panel does not capture the full diversity of MCBAs present *in vivo*. BA concentrations also vary with factors such as diet and circadian rhythms, and the concentrations encountered by *C. difficile* in the gut may differ from those examined *in vitro*. Studies have found varying concentrations of MCBAs *in vivo,* and although the concentrations used in this paper are physiologically relevant, in a host these concentrations are not static (21, 22) (44, 45). We also only used one strain of *C. difficile* in this study, although it is a clinically relevant strain R20291. Different strains might have different responses to these MCBAs. Finally, we focused on *in vitro* assays in a rich BHI medium versus a defined medium or an *ex vivo* assay.

Future work will focus on how BSHs are able to modulate amino acid concentrations in the gut through their reconjugation and deconjugation activity of MCBAs. Do BSHs make or break MCBAs to sequester amino acids from the gut, and or liberate amino acids for support *C. difficile* growth? It is known that *C. difficile* competes with other members of the gut microbiota for amino acids and this competition for nutrients reduces *C. difficile* colonization and pathogenesis (7, 9). However, recent studies show that nutrient limitation alone may not be sufficient in preventing *C. difficile* growth, and that secondary BA synthesis may be needed to reduce virulence (46). What is not known is how MCBAs influence this process. Further analysis in defined media supplemented with amidates of those auxotrophies is needed to determine if *C. difficile* can access amino acids conjugated to BAs, and if BSHs can reconjugate amino acids to reduce key amino acids *in vitro*.

Ultimately, our findings demonstrate that MCBAs are functionally diverse and that their effects cannot be predicted solely from conjugation status or amino acid biochemical properties. Specific MCBAs may warrant further investigation as potential targets for BSH-based or MCBA-based therapeutic strategies. Although their contributions to CR and disease *in vivo* remain to be established, MCBAs represent an important new metabolite class for understanding microbiota-mediated protection against CDI and other intestinal diseases.

## Acknowledgements

C.M.T. Research reported in this publication was supported by the National Institute of General Medical Sciences of the National Institutes of Health under award number R35GM149222 and the National Institute of Diabetes and Digestive and Kidney Diseases of the National Institutes of Health under award number R01DK148700. The content is solely the responsibility of the authors and does not necessarily represent the official views of the National Institutes of Health. S.W.H is funded by the US Army Medical Department and the Long-term Health Education and Training Scholarship.

## Author Contributions

Conceptualization, S.W.H and C.M.T; Data Curation, S.W.H. and N.S.L.; Formal Analysis, S.W.H., B.C. and N.K.N.; Funding acquisition, E.C.G and C.M.T.; Investigation, S.W.H. and C.E.P.; Methodology, S.W.H., N.S.L. and N.K.N.; Project Management E.C.G. and C.M.T.; Resources, E.C.G. and C.M.T.; Software, S.W.H.; Supervision, E.C.G. and C.M.T; Validation, S.W.H.; Visualization, S.W.H. and C.M.T.; Writing-Original Draft, S.W.H.; Writing-review and editing, N.S.L., A.S.G., S.W.H. and C.M.T.

## Declaration of generative AI and AI-assisted technologies in the writing process

No Generative AI was not used in the writing process.

## Supplemental Figure legends

**Figure S1: CA amidates and *C. difficile* growth kinetics, spores, and toxin activity over time.**

A-D) *C. difficile* growth curves measuring OD_600_ over 24 hr in varying concentrations of AA-CAs. Concentrations ranged from 0 mM (2.5% DMSO), 1.25 mM, 2.5 mM, 5 mM, and 10 mM. Conditions were run in triplicate with three technical replicates. E) MIC assay of Tyr-CA after incubation in differing concentrations of Tyr-CA for 24hr. Concentrations ranged from 1.25 mM, 2.5 mM, 5 mM, and 10 mM. Inhibition was determined if after 24 hr growth did not exceed inoculum (dotted line). F) Enumeration of vegetative cells (solid bars) and spores (striped bars) after incubation in 2.5 mM MCBAs at 24 and G) 48 hr. Significance was determined by ordinary one-way ANOVA with Dunnett’s posttest comparing each MCBA to CA. H) Toxin activity via a Vero Cell cytotoxicity assay after incubation in 2.5 mM AA-CAs for 72 hr, significance was determined by ordinary one-way ANOVA with Dunnett’s posttest comparing each MCBA to CA. * = p< 0.05, ** = p<0.01, ***= p<0.001, **** = p< 0.0001.

**Figure S2: DCA amidates and *C. difficile* growth kinetics, spores, and toxin activity over time.** A-D) *C. difficile* growth curves measuring OD_600_ over 24 hr in varying concentrations of AA-DCAs. Concentrations ranged from 0 mM (2.5% DMSO), 0.075 mM, 0.3 mM, 0.6 mM, and 1.2 mM. Conditions were run in triplicate with three technical replicates. E) Enumeration of vegetative cells (solid bars) and spores (striped bars) after incubation in 0.3 mM AA-DCAs for 24 and F) 48 hr. Significance was determined by ordinary one-way ANOVA with Dunnett’s posttest comparing each MCBA to DCA G) Toxin activity via a Vero Cell cytotoxicity assay after incubation in 0.3 mM AA-DCAs for 24 hr Significance was determined by ordinary one-way ANOVA with Dunnett’s posttest comparing each MCBA to DCA. * = p< 0.05, ** = p<0.01, ***= p<0.001, **** = p< 0.0001.

**Figure S3: CDCA amidates and *C. difficile* growth kinetics, spores, and toxin activity over time.** A-D) *C. difficile* growth curves measuring OD_600_ over 24 hr in varying concentrations of AA-CDCAs. Concentrations ranged from 0 mM (2.5% DMSO), 0.075 mM, 0.3 mM, 0.6 mM, and 1.2 mM. Conditions were run in triplicate with three technical replicates. E) Enumeration of vegetative cells (solid bars) and spores (striped bars) after incubation in 0.3 mM AA-CDCAs for 24 and F) 48 hr. Significance was determined by ordinary one-way ANOVA with Dunnett’s posttest comparing each MCBA to CDCA G) Toxin activity via a Vero Cell cytotoxicity assay after incubation in 0.3 mM AA-CDCAs for 24 hr Significance was determined by ordinary one-way ANOVA with Dunnett’s posttest comparing each MCBA to CDCA. * = p< 0.05, ** = p<0.01, ***= p<0.001, **** = p< 0.0001.

**Figure S4: LCA amidates and *C. difficile* spores and toxin activity over time.** A-B) Enumeration of vegetative cells (solid bars) and spores (striped bars) after incubation in 0.25x MIC AA-LCAs for 24 and 48 hr. Significance was determined by ordinary one-way ANOVA with Dunnett’s posttest comparing each MCBA to LCA. C) Toxin activity via a Vero Cell cytotoxicity assay after incubation in 0.25x MIC mM AA-LCAs for 24 hr Significance was determined by ordinary one-way ANOVA with Dunnett’s posttest comparing each MCBA to LCA. * = p< 0.05, ** = p<0.01, ***= p<0.001, **** = p< 0.0001.

## Materials and Methods

### Bacterial cultures

*C. difficile* R20291 spores were plated on Brain Heart Infusion (BHI) Agar supplemented with 10mg/L L-cysteine, and 10% (wt/vol) Taurocholate and incubated at 37°C overnight in an anaerobic chamber (Coy Labs). One colony from the germinated spores was inoculated into 5mL of BHI broth+ 10 mg/mL L-cysteine and incubated at 37°C for 16-18 hours. After the incubation, the culture was diluted 1:5 in fresh BHI broth+ cysteine, and the OD_600_ was measured. The culture was allowed to grow for about 4 hours until the OD_600_ doubled to allow the culture to reach midlog. The resulting cultures were diluted to the appropriate OD_600_ for desired experimental use.

### Spore preparation

Spore preparations were prepared as previously described (Theriot et al, 2014). Briefly, *C. difficile* strain R20291 was incubated for 7 days in CloSpore media. After 7 days, cultures were centrifuged, washed with water, and then resuspended in 1mL of sterile water. Resulting stocks were heat treated at 65°C for 20 minutes to kill vegetative cells, and plated on TBHI plates to enumerate the amount of spores/mL. Spores were stored at 4°C in between uses. **MCBA synthesis.** Organic solutions were concentrated under reduced pressure on a Büchi rotary evaporator using a water bath. Chromatographic purification of products was accomplished by flash chromatography on Silicycle F60 silica gel. All reactions were carried out in well ventilated fume hoods. Thin-layer chromatography (TLC) was performed on Silicycle 250 μm silica gel plates. Visualization of the developed chromatogram was performed by irradiation with 254 nm UV light or treatment with a solution of ceric ammonium molybdate stain followed by heating. Yields refer to purified compounds unless otherwise noted. ^1^H and ^13^C NMR spectra were recorded on a Bruker Advance IV Neo 400 MHz, 500 MHz, 600 MHz and Bruker Advance III 700 MHz instruments and are internally referenced to residual protiosolvent signals of CD_3_OD at δ 3.31 and 49.0 ppm for ^1^H and ^13^C, respectively. Data for ^1^H NMR are reported as follows: chemical shift (δ ppm), integration, multiplicity (s = singlet, br s = broad singlet, d = doublet, t = triplet, q = quartet, m = multiplet), and coupling constant (Hz). Data for ^13^C NMR are reported in terms of chemical shift and no special nomenclature is used for equivalent carbons. MCBA synthesis was adapted from a previously reported protocol (Ezawa et. al, 2017). In our protocol, the bile acid core (0.25 mmol, 1 eq.) was dissolved in anhydrous THF (4.9 ml, 0.05 M) and allowed to stir at 0 °C. Ethyl chloroformate (0.3 mmol, 1.2 eq.) was then added, followed by triethylamine (0.3 mmol, 1.2 eq.) and the reaction was stirred at 0 °C for 1-2 h. After complete conversion of starting material by TLC, a cold solution of amino acid (0.375 mmol, 1.5 eq.) and inorganic base (NaHCO_3_ or NaOH, 1.5 eq.) in H_2_O (4.9 ml, 0.05 M) was added in one portion at 0 °C. The reaction was stirred for 2 h, gradually allowing it to warm to room temperature. After this time, the crude reaction mixture was concentrated under vacuum to remove THF, then 2 M HCl was added to acidify the aqueous mixture to pH < 2, producing a white precipitate. The mixture was extracted in dichloromethane (3 × 20 ml) and the combined organic layers were washed with brine (50 ml), dried over Na_2_SO_4_ and concentrated *in vacuo*. The crude residue was purified over silica gel using a mixture of methanol and dichloromethane as described below to produce the desired products. All other structures were synthesized according to literature procedure, and characterization data is consistent with reported data.

### Growth kinetics

#### Plate reader assay

Exponentially growing *C. difficile* cells were diluted to an OD_600_ of 0.2 in 1mL of BHI broth. 10µL of the culture was used to inoculate 190µL of varying MCBA concentrations on a clear, flat bottomed 96-well plate (Genesee scientific) for a resulting starting OD_600_ of 0.1 in each well. MCBA conditions started at the recorded MIC of the sterol core and halved until either 1/8^th^ the MIC or 1/16^th^ the MIC. Conditions were made by adding stocks of MCBAs dissolved in DMSO into 1.5 mL of BHI broth, then aliquoted into the 96 well plate. MCBA conditions other than the MIC condition had additional DMSO added to a total of 2.5% (v/v) to standardize vehicle concentrations across all conditions. *C. difficile* in BHI alone and in 2.5% DMSO were used as controls. Plates were run in a TECAN sunrise plate reader and anaerobically at 37°C with shaking for 24hr, and OD_600_ measurements taken each hour. Each condition was run in triplicate, and three biological replicates were performed.

### Propidium Iodide (PI) Stain assay

4mL of exponentially grown *C. difficile* cells were spun down, washed with PBS twice, and then combined and resuspended in PBS. Conditions of bile acids at 1/4^th^ the MIC of the sterol core were inoculated to have a final OD_600_ of 0.1 and incubated at 37°C for 30 minutes. A 150µM SDS condition was used as a positive control for membrane disruption. Following the incubation, cells were stained with Propidium Iodide at 20µg/mL for 30 min at 37°C. Stained bacteria were diluted 1:10 in PBS and excitation was read on a clear, flat bottomed 96-well plate (absorbance: 540nm, excitation 610nm) in a Tecan Infinite F200 pro plate reader and background fluorescence was subtracted.

### MIC Assay

MIC assay was modified from the MIC assay performed in (Foley et al 2023). Exponentially growing *C. difficile* cells were back-diluted to a starting OD_600_ of 0.01 and incubated for 24 h at 37°C. Drip plating was performed at 0 h to determine the inoculum, and again at 24 h to measure overall growth. The MIC was determined as the lowest concentration that inhibited a doubling from the starting inoculum after 24 h.

### Sporulation Assay

Exponentially growing *C. difficile* cells were inoculated into BHI media containing 0.25xMIC at an OD_600_ of 0.01 and incubated at 37°C for 72 hours. Every 24 hours, samples were taken, enumerated on BHI agar plates for the presence of vegetative cells, and then heated at 60°C for 20 minutes and plated on TBHI for enumeration of spores only.

### Vero Cell Cytotoxicity Assay

Toxin activity was measured using a Vero cell cytotoxicity assay. Vero cells are grown and maintained in DMEM media (Gibco Laboratories, 11965-092) with 10% fetal bovine serum (Gibco Laboratories, 16140-071) and 1% penicillin-streptomycin solution (Gibco Laboratories, 15070-063). Cells were incubated with 0.25% trypsin (Gibco Laboratories, 25200-056), washed with 1x DMEM media, and harvested by centrifugation at 250 x g for 5 min. Cells were seeded at 1 x 10^-4^ cells per well in a 96-well flat-bottom microtiter plate and incubated overnight at 37°C/5% CO_2_. Media samples were only filtered before testing. The samples were further diluted 10-fold to a maximum of 10^-6^ Sample dilutions were incubated 1:1 with PBS (for all dilutions) or antitoxin (TechLab, T5003) for 40 min at room temperature. Following the incubation, these mixtures were added to the Vero cells and plates were incubated overnight at 37°C/5% CO_2_. Vero cells were viewed under 200X magnification for rounding after overnight incubation. The cytotoxic titer was defined as the reciprocal of the highest dilution that produced rounding in 80% of Vero cells for each sample. Vero cells treated with purified *C. difficile* toxin A were used as controls.

### Statistical analysis

Statistical analysis was performed using Prism 10 for macOS (GraphPad Software, La Jolla, CA, USA). For the growth rate analysis, PI Stain, Toxin, and Sporulation, significance was determined through a one-way analysis of variance (ANOVA) with Dunnets multiple comparisons correction. All tests were run with set statistical significance of p<0.05 and at least three biological replicates.

