## Supplementary material for "Microbially conjugated bile acids differentially affect the lifecycle and pathogenesis of *Clostridioides difficile*": Figure S1

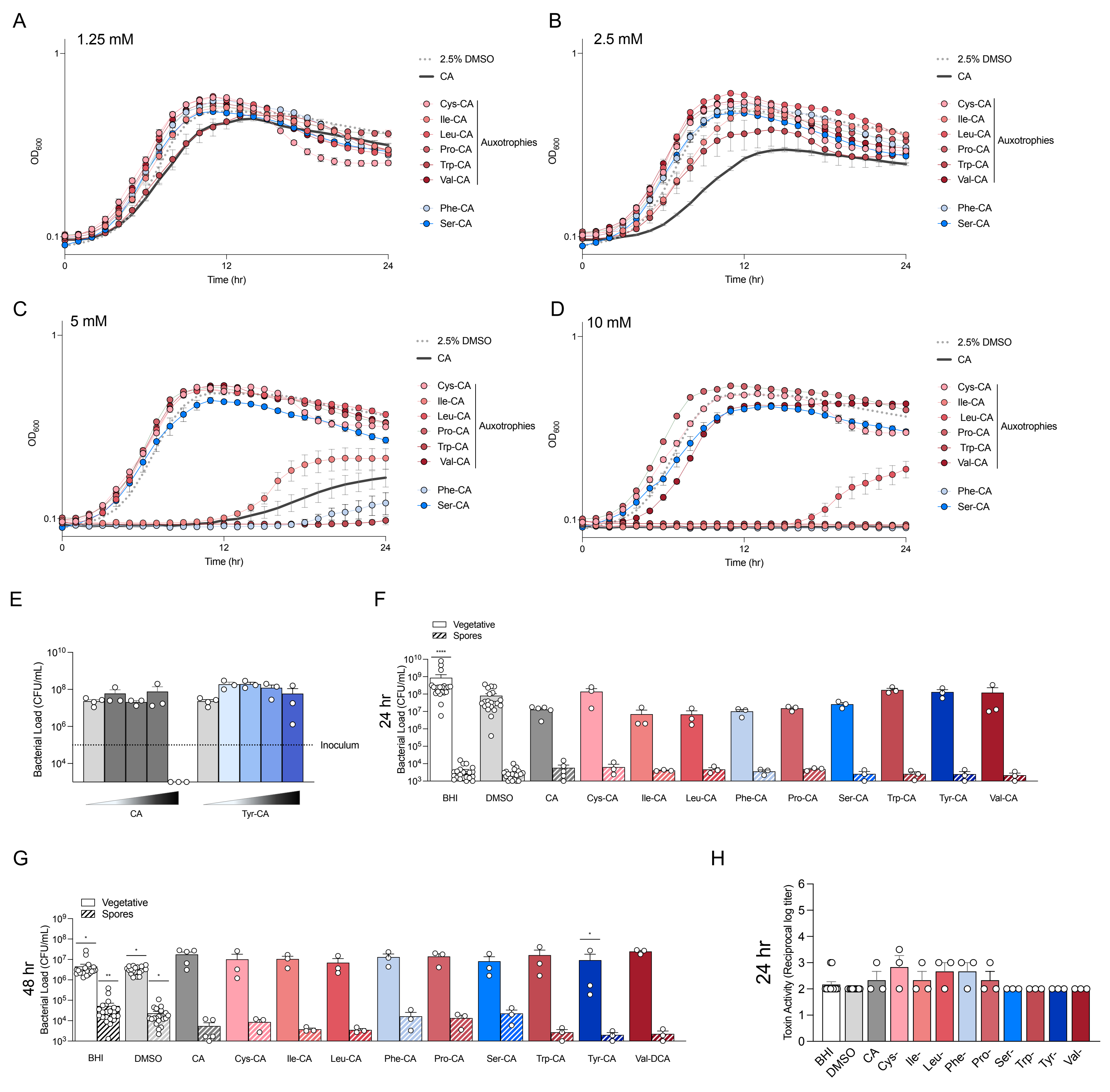

A
1.25 mM
B
2.5 mM
C
5 mM
D
10 mM
OD600
Time (hr)
2.5% DMSO
CA
Cys-CA
Ile-CA
Leu-CA
Pro-CA
Trp-CA
Val-CA
Auxotrophies
Phe-CA
Ser-CA
E
Bacterial Load (CFU/mL)
Inoculum
CA
Tyr-CA
F
Vegetative
Spores
24 hr
BHI
DMSO
Tyr-CA
Val-CA
G
48 hr
Val-DCA
H
Toxin Activity (Reciprocal log titer)
