## Supplementary material for "Microbially conjugated bile acids differentially affect the lifecycle and pathogenesis of *Clostridioides difficile*": Figure S2

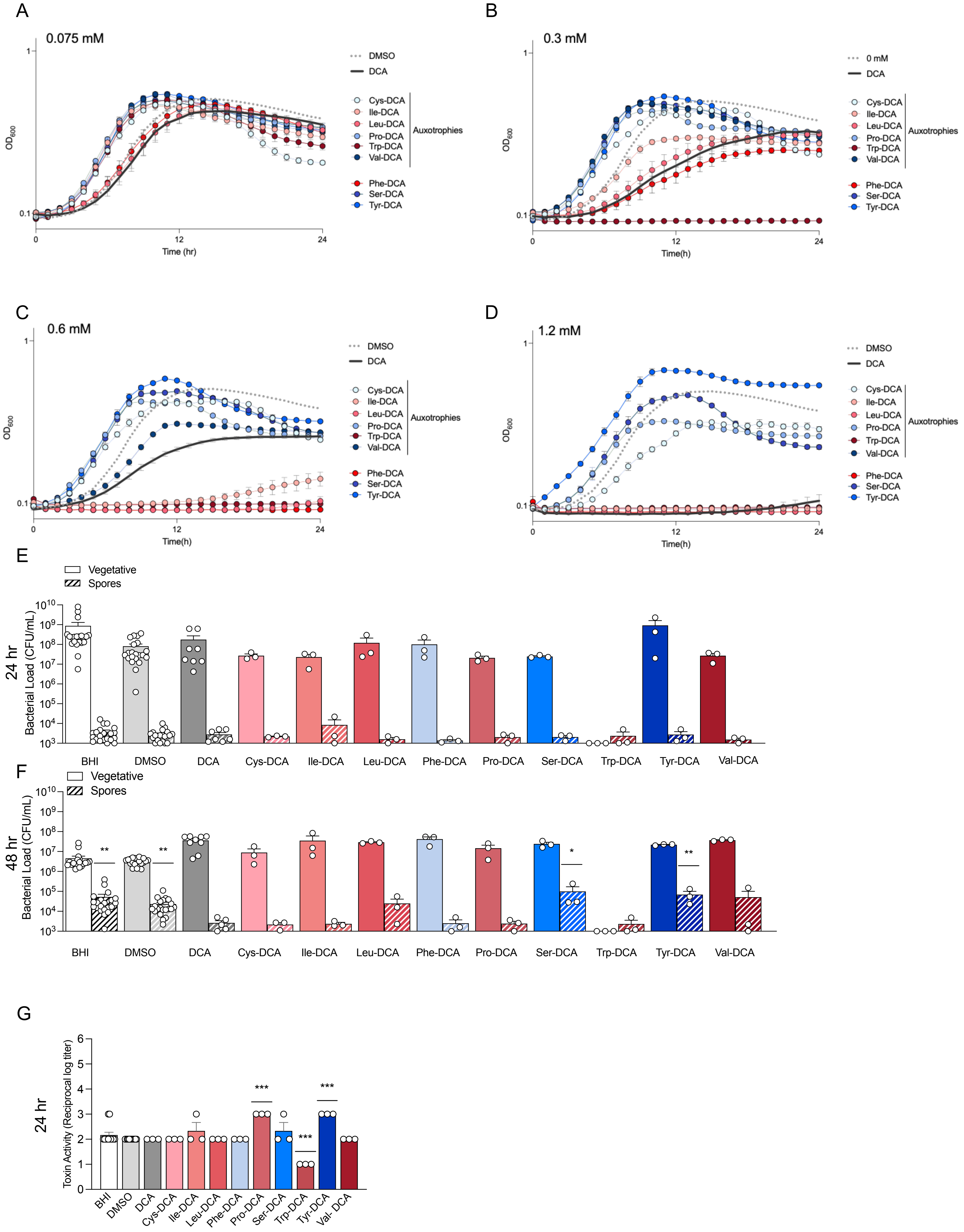

A
0.075 mM
OD600
Time (hr)
DMSO
DCA
Cys-DCA
Ile-DCA
Leu-DCA
Pro-DCA
Trp-DCA
Val-DCA
Auxotrophies
Phe-DCA
Ser-DCA
Tyr-DCA
B
0.3 mM
Time(h)
0 mM
C
0.6 mM
D
1.2 mM
E
Vegetative
Spores
24 hr
Bacterial Load (CFU/mL)
BHI
F
48 hr
G
Toxin Activity (Reciprocal log titer)
Val- DCA
