## Supplementary material for "Microbially conjugated bile acids differentially affect the lifecycle and pathogenesis of *Clostridioides difficile*": Figure S3

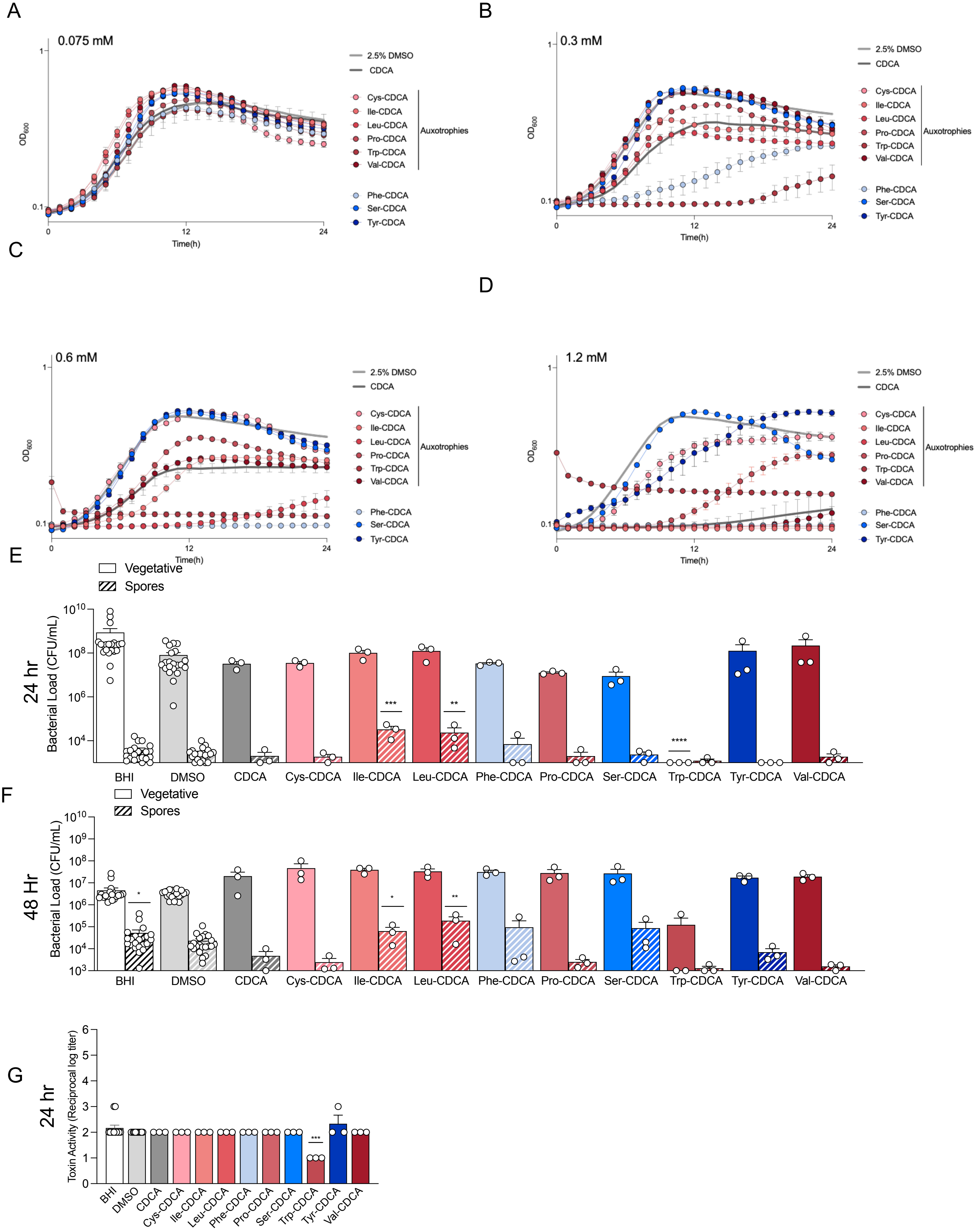
A
0.075 mM
OD600
Time(h)
2.5% DMSO
CDCA
Cys-CDCA
Ile-CDCA
Leu-CDCA
Pro-CDCA
Trp-CDCA
Val-CDCA
Auxotrophies
Phe-CDCA
Ser-CDCA
Tyr-CDCA
B
0.3 mM
C
0.6 mM
D
1.2 mM
E
Vegetative
Spores
24 hr
Bacterial Load (CFU/mL)
BHI
DMSO
CDCA
Cys-CDCA
Ile-CDCA
Leu-CDCA
Phe-CDCA
Pro-CDCA
Ser-CDCA
Trp-CDCA
Tyr-CDCA
Val-CDCA
F
48 Hr
G
Toxin Activity (Reciprocal log titer)
