## Supplementary material for "Microbially conjugated bile acids differentially affect the lifecycle and pathogenesis of *Clostridioides difficile*": Figure S4

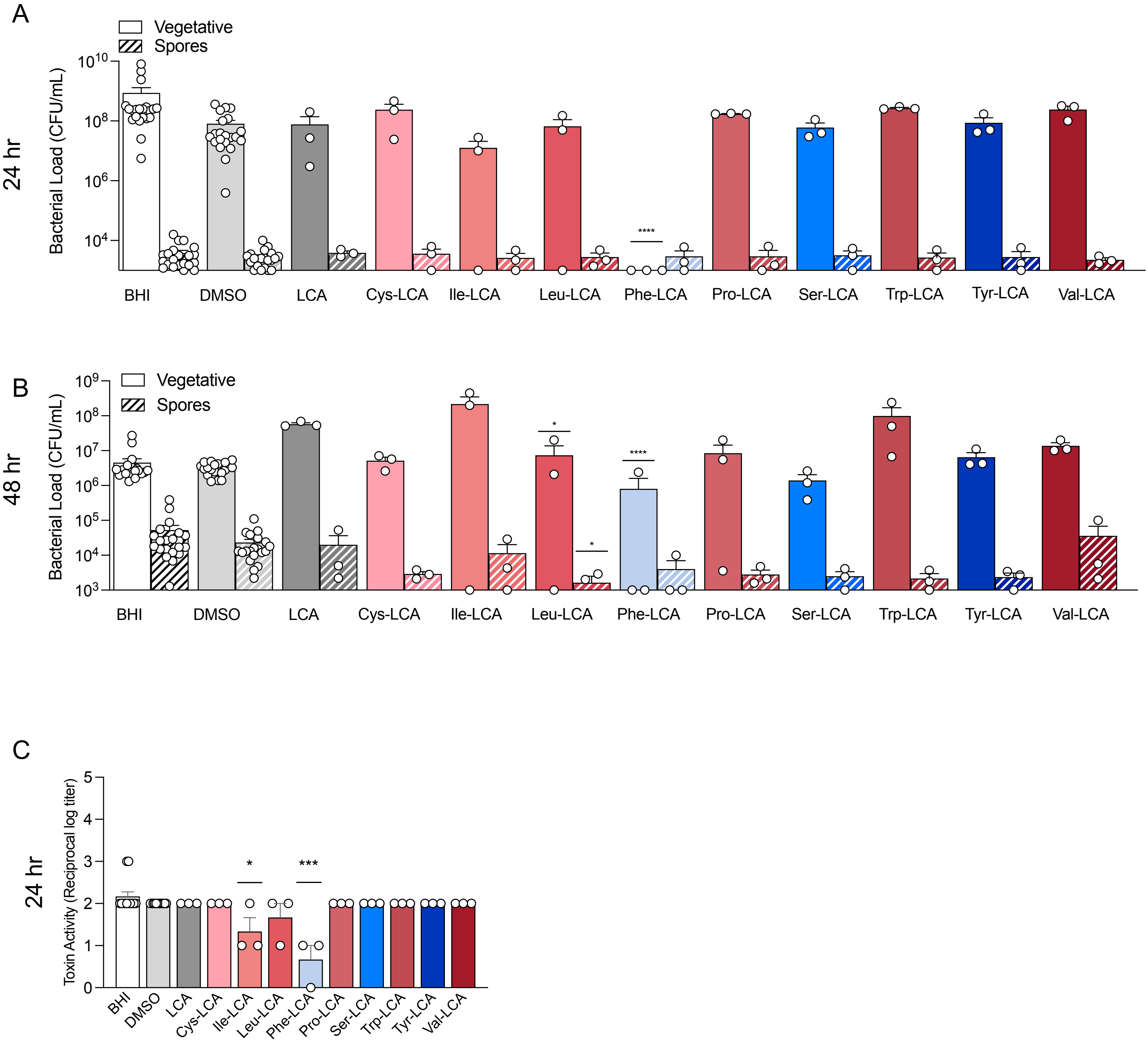

A
Vegetative
Spores
24 hr
Bacterial Load (CFU/mL)
$10^{10}$
$10^{8}$
$10^{6}$
$10^{4}$
****
BHI
DMSO
LCA
Cys-LCA
Ile-LCA
Leu-LCA
Phe-LCA
Pro-LCA
Ser-LCA
Trp-LCA
Tyr-LCA
Val-LCA
B
Vegetative
Spores
48 hr
Bacterial Load (CFU/mL)
$10^{9}$
$10^{8}$
$10^{7}$
$10^{6}$
$10^{5}$
$10^{4}$
$10^{3}$
*
*
****
BHI
DMSO
LCA
Cys-LCA
Ile-LCA
Leu-LCA
Phe-LCA
Pro-LCA
Ser-LCA
Trp-LCA
Tyr-LCA
Val-LCA
C
24 hr
Toxin Activity (Reciprocal log titer)
5
4
3
2
1
0
*
***
BHI
DMSO
LCA
Cys-LCA
Ile-LCA
Leu-LCA
Phe-LCA
Pro-LCA
Ser-LCA
Trp-LCA
Tyr-LCA
Val-LCA
